# A systematic evaluation of SIRT6 as a transcriptomic biomarker of aging

**DOI:** 10.64898/2026.09.03.749075

**Authors:** E. Kashuk, A. Tarakanova, A. Malygina, N. Kuzovkina, A. Ponomareva, D. Toiber, E. Khrameeva, D. Smirnov

## Abstract

SIRT6 is a NAD^+^-dependent sirtuin that plays central roles in chromatin regulation, DNA repair, telomere maintenance, metabolic homeostasis, and inflammatory control. Although SIRT6 have long been implicated in aging because of its association with several hallmarks of aging, the available evidence remains largely context-dependent and mechanistic, limiting the interpretation of SIRT6 as a robust and evolutionarily conserved biomarker of aging. To comprehensively investigate the role of SIRT6 as an aging biomarker, we established SIRT6.db, a multi-species transcriptomic resource that integrates SIRT6-targeted perturbation experiments across diverse biological systems and organisms form all available SIRT6-related publications collected via a large-scale textual analysis of the SIRT6 literature using topic modeling. Based on this database, we identified both species-specific and evolutionarily conserved transcriptional and functional signatures associated with SIRT6 perturbation and established their relevance to hallmarks of aging. We showed that SIRT6 expression is generally stable during normal aging, but becomes dysregulated in Alzheimer’s disease in cell type and stage-specific manner, highlighting the context dependence of its potential as a transcriptomic biomarker of aging.

## 1 Introduction

Aging is a multifactorial biological process characterized by the progressive decline of physiological integrity, increased vulnerability to disease, and ultimately reduced survival. During the past decades, substantial effort has been devoted to identifying conserved molecular mechanisms that control aging and longevity across species. This effort has been advanced by introducing the hallmarks of aging [1, 2], that organize age-associated processes into interconnected features such as genomic instability, telomere attrition, epigenetic alterations, loss of proteostasis, mitochondrial dysfunction, cellular senescence, and altered intercellular communication. Within this conceptual framework, aging biomarkers have emerged as a critical tool for quantifying biological age, predicting functional decline and evaluating responses to geroprotective interventions [3, 4]. Such biomarkers are typically identified through integrative analyses of molecular, cellular, and physiological data, including transcriptomic, epigenomic, proteomic, and clinical measurements [3, 5–9]. However, their utility depends on robust validation across tissues, experimental systems, and longitudinal settings to demonstrate reproducibility, specificity and predictive value [6, 10, 11].

The role of the sirtuin family (SIRT1-SIRT7) in aging has been extensively investigated over the past several decades. Sirtuins are NAD^+^-dependent enzymes characterized by distinct histone deacetylase activities and specific subcellular localizations. Previous studies have shown that sirtuins are involved in the regulation of genome stability [12, 13], maintenance of telomere integrity [14, 15], control of cellular metabolism [16–19], and modulation of gene expression [13, 20, 21], supporting their consideration as promising candidates for aging biomarkers. For example, SIRT1 was linked to age-associated metabolic decline and inflammatory signaling [22–24], and its activity was associated with mechanisms underlying caloric restriction [25–27]. SIRT3, a major mitochondrial sirtuin, was also associated with impaired oxidative metabolism, oxidative stress, and age-related functional decline across tissues [28]. Additionally, SIRT6 has ben shown as a particularly compelling candidate biomarker because of its established roles in chromatin regulation, DNA repair and telomere maintenance, all of which are central to organismal aging [29–33].

At the same time, the role of sirtuins as evolutionarily conserved regulators of longevity has been questioned. Lifespan extension initially reported in model organisms was later challenged by studies in worms and flies, which attributed the observed effects to confounding factors, including a secondary mutation in one of the *sir-2.1* transgenic *C. elegans* strains and GAL4 driver effects in flies [34, 35]. A subsequent study further showed that the effect of *sir-2.1* overexpression on *C. elegans* lifespan was influenced by the thymidylate synthase inhibitor 5-fluorodeoxyuridine (FUDR), which is commonly used to prevent progeny development [36]. In vertebrates, SIRT6 has been the sirtuin most consistently linked to longevity [37, 38]. However, this effect appears to be sex-specific [39] and remains far from being established as species-conserved [40]. Moreover, SIRT6 still lacks robust predictive validation using cross-sectional and longitudinal multi-omics datasets, which is essential for assessing its utility as a reliable biomarker of aging.

Despite the substantial body of accumulated evidence, the causal mechanisms linking sirtuin activity to aging processes remain insufficiently understood, and their applicability as universal biomarkers of aging has not been convincingly established. The present study aims to provide a clear overview of the role of the sirtuin family in aging, with particular focus on one of its most promising members, SIRT6. Using literature-scale topic modeling and a multi-species transcriptomic resource, we show that SIRT6 is linked to several aging hallmarks, but that these associations are dynamic, context-dependent, and only partially conserved across biological systems. We further examine SIRT6 expression in human aging and Alzheimer’s disease (AD) datasets, revealing tissue-, cell-type-, and disease-specific patterns that refine the interpretation of SIRT6 as a candidate biomarker of aging.

## 2 Results

### 2.1 Topic modeling reveals hallmark-specific themes in SIRT6 research

To achieve a comprehensive and systematic assessment of the aging-related functions of SIRT6 based on the existing literature, we conducted a textual analysis of all publicly available articles on SIRT6 using a topic modeling approach (Fig. 1a). Specifically, we compiled a corpus of 1648 abstracts from SIRT6-related research articles and preprints retrieved from the OpenAlex [41] and Europe PMC [42] databases by querying for SIRT6 in the abstract text. To confirm the relevance of the selected abstracts to SIRT6, we performed an additional centrality analysis and retained only papers with a centrality score greater than 0.4, thereby excluding records in which SIRT6 was mentioned only incidentally (Supplementary Fig. 1a). A detailed description of the corpus construction procedure is provided in the Methods section.

**Fig. 1.**
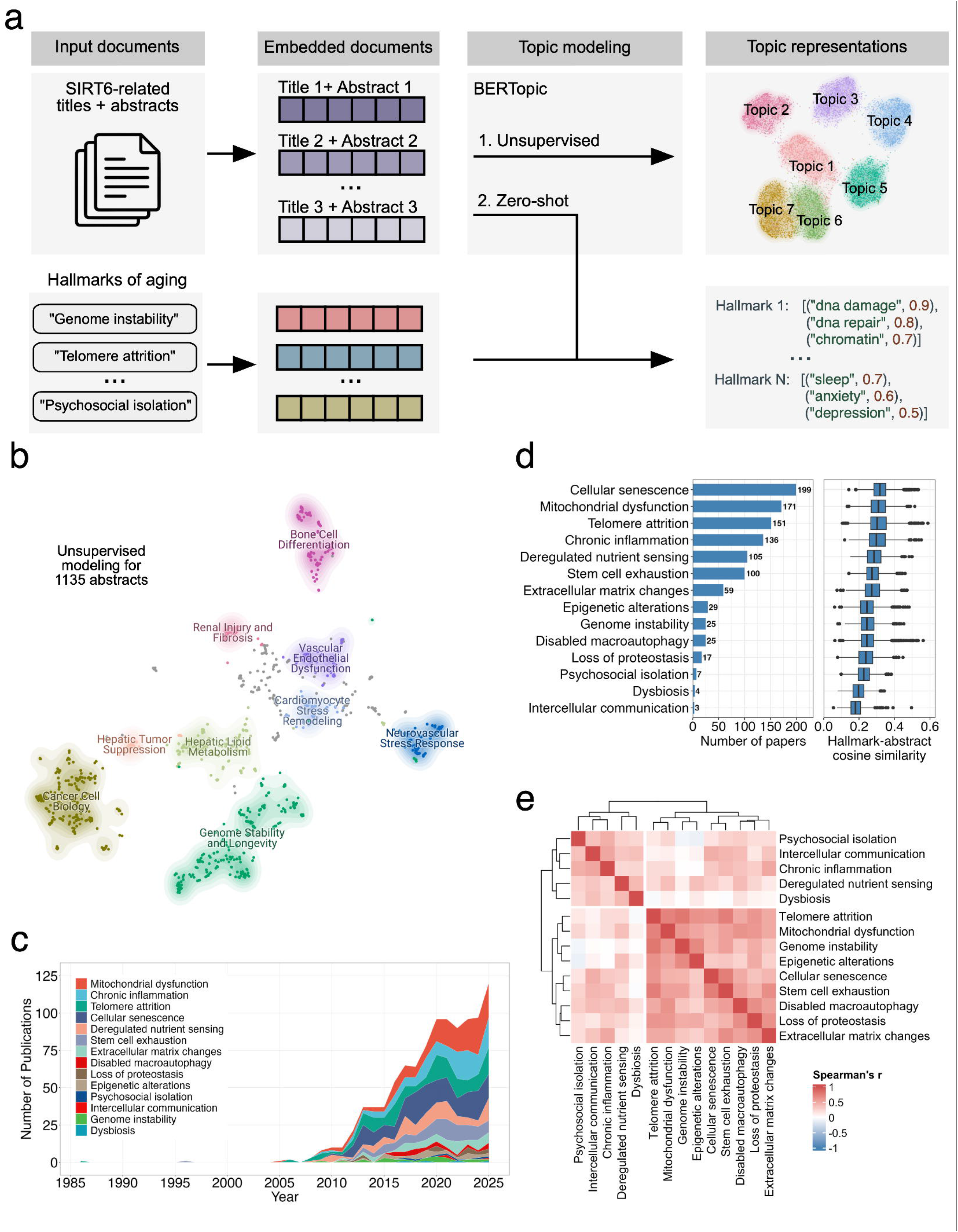
Topic modeling analysis of SIRT6-related research articles. (a) Overview of the analysis pipeline. Combined titles and abstracts from SIRT6-related articles and preprints were used as input for BERTopic-based topic modeling in unsupervised and zero-shot modes. Zero-shot analysis was performed using a predefined list of topic labels corresponding to the hallmarks of aging. (b) UMAP visualization of the resulting document clusters, labeled according to the aggregated keywords for each topic. (c) Stacked area plot showing the number of articles associated with each hallmark over time. (d) Distribution of articles classified under each aging hallmark (left) and distribution of article-level probabilities of association with aging hallmarks (right). (e) Pairwise Spearman correlation matrix for hallmark-related terms identified in the zero-shot analysis.

Unsupervised analysis of the collected corpus of previously published studies on SIRT6 (Supplementary Table 1) identified two major semantic clusters corresponding to investigations of its role in oncogenesis (“Cancer Cell Biology”, 225 documents) and in aging-related processes (“Genome Stability and Longevity”, 223 documents) (Fig. 1b, Supplementary Table 2). The remaining identified clusters were predominantly tissue- or organ-specific, highlighting the diversity of biological systems in which SIRT6 has been investigated.

To further establish the association of SIRT6-related papers with specific hallmarks of aging, we performed zero-shot topic modeling using hallmark definitions as predefined topic terms (Supplementary Table 3). The number of SIRT6-related publications focusing on aging hallmarks increased steadily across publication years, indicating growing interest in the aging-related functions of SIRT6 (Fig. 1c). However, the relative focus of this literature also shifted over time. Cellular senescence, which has traditionally been one of the most abundant hallmark categories in SIRT6-related research, has recently been complemented by increasing attention to mitochondrial dysfunction and chronic inflammation hallmarks, changing focus toward metabolic and inflammatory dimensions of aging. Overall, our analysis revealed that SIRT6 functions were most strongly associated with cellular senescence (199 documents), mitochondrial dysfunction (171 documents) and telomere attrition (136 documents) (Fig. 1d). Interestingly, loss of proteostasis remains one of the least studied hallmarks of aging in the context of SIRT6 activity, despite recent evidence suggesting that proteostasis may be one of the key regulators of aging-related phenotypes [43]. These associations were also found to display pronounced tissue- and organ-specific patterns. In particular, the link between SIRT6 and mitochondrial dysfunction was observed predominantly in heart and skeletal muscle models.

Because most hallmarks of aging are known to be interconnected, we next estimated relationships between hallmark categories in our corpus using abstract-hallmark cosine similarity values. Specifically, pairwise Spearman correlations were calculated across hallmark similarity profiles, followed by hierarchical clustering of the resulting correlation matrix (Fig. 1e). This analysis separated the hallmarks into two groups. The first group corresponded primarily to processes that occur within cells or in their immediate tissue microenvironment (e.g. genome instability, telomere attrition, epigenetic alterations, cellular senescence). The second group mostly comprised integrative hallmarks, associated with a tissue homeostasis (deregulated nutrient sensing, altered intercellular communication, chronic inflammation, dysbiosis, and psychosocial isolation). Additionally, we performed hallmark co-occurance analysis, which revealed a tightly connected module of telomere attrition, cellular senescence and stem cell exhaustion hallmarks, with the strongest positive three-hallmark associations (lift scores *>* 5.83) (Supplementary Table 4). Together, these results implicate SIRT6 in both initiating mechanisms and downstream consequences of aging.

### 2.2 Cross-species public transcriptomic data reveal aging-related effects of SIRT6 perturbation

Although prior experimental studies strongly implicate SIRT6 in most of the aging hallmarks, most evidence are derived from selected models, tissues, or perturbation experiments. Therefore we were wondering if its relevance to mechanisms of aging is detectable and conserved across diverse biological systems. To answer this question, we took an advantage of publicely available bulk RNA-seq datasets on SIRT6 perturbations, as the most abundant type of omics data available, to construct a comprehensive cross-species transcriptomics database of SIRT6-targeted experiments. SIRT6-targeted studies from six organisms (*Homo sapiens*, *Mus musculus*, *Rattus norvegicus*, *Macaca fascicularis*, *Sus scrofa*, and *Drosophila melanogaster* ) were included (Fig. 2a). The resulting resource, SIRT6.db, is publicly available at https://sirt6.github.io/SIRT6.db/. Additionally, we classify the experiments into nine biological systems (nervous, cardiovascular, developmental, cancer, respiratory, metabolic, gastrointestinal, musculoskeletal, and immune).

**Fig. 2.**
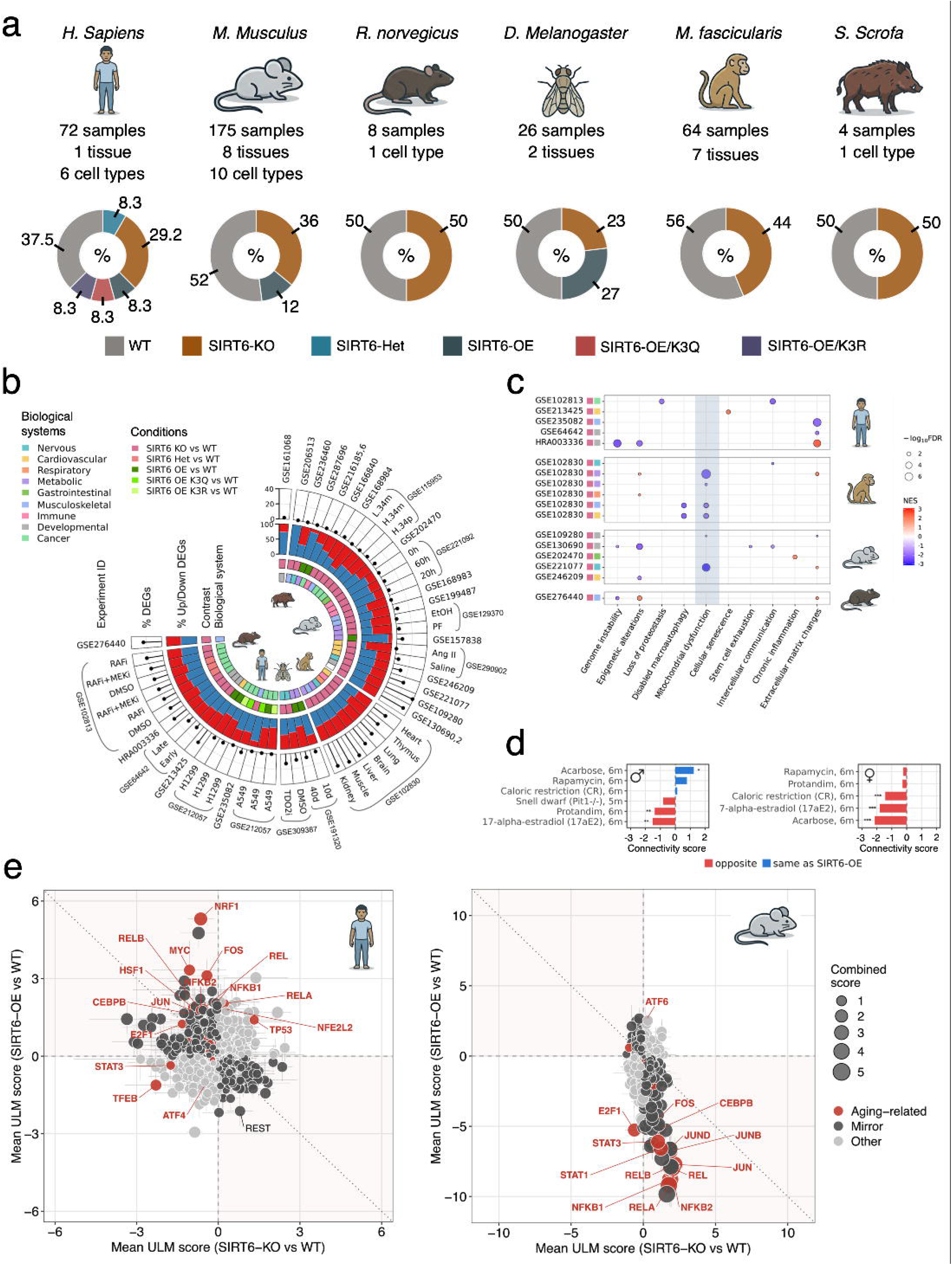
Transcriptomic landscape of SIRT6-targeted models across species. (a) Overview of the assembled cross-species transcriptomics datasets for SIRT6-targeted studies, demonstrating the percentage of each biological condition (WT, SIRT6-KO, SIRT6-OE, etc) and the total number of experimental replicates across the species. (b) The summary of the DE analysis that shows the magnitude and composition of transcriptional changes across organisms, biological systems and contrasts. Red bars correspond to the proportion of upregulated genes, while blue bars shows the percentages of downregulated genes across experiments. (c) Gene set enrichment analysis of aging hallmarks across SIRT6-KO versus WT contrasts from six species. (d) Sex-specific associations between transcriptomic signatures of longevity interventions and hepatic SIRT6-OE. (e) Analysis of the activity of TFs implicated into the regulation of DE genes in SIRT6-KO and SIRT6-OE models in human (left panel) and mouse (right panel).

First, we estimated the magnitude of expression changes between SIRT6-targeted samples and their corresponding controls. Differential expression analysis revealed that SIRT6 alterations induce measurable transcriptional changes across all 54 analyzed datasets (Fig. 2b). However, the percentage of differentially expressed genes (DEGs) varied between experiments, with some datasets showing more pronounced transcriptional alterations (macaca, human and rat), while others exhibit smaller changes (mouse, hog and fly). Although the proportion of DEGs was generally low across mouse datasets, developmental cell models (e.g. embryonic fibroblasts and stem cells) exhibited the most extensive transcriptional alterations following SIRT6 deficiency (Fig. 2b). Also, we observed no significant and consistent bias in gene expression changes towards either shorter or longer genes across species (Supplementary Fig. 2a). Thus, SIRT6 does not have a uniform transcriptional effect, it rather regulates in a context-dependent manner.

To determine whether transcriptional consequences of SIRT6 loss are associated with canonical aging hallmarks, we performed gene set enrichment analysis on SIRT6-KO versus WT samples (Fig. 2c) using hallmark gene signatures from Open Genes database [44]. Significant enrichment (FDR-adjusted *p*-value *<* 0.05) was detected for 9 out of the 14 canonical hallmarks. Although epigenetic alterations and extracellular matrix changes were the most frequently enriched hallmarks, the direction of enrichment varied across species. Mitochondrial dysfunction was the most consistently enriched hallmark: across independent experiments in *Homo sapiens*, *Macaca fascicularis*, and *Mus musculus*, the normalized enrichment score was negative in every case, indicating consistent downregulation of mitochondrial gene programs upon SIRT6 knockout. An independent enrichment analysis using MSigDB Hallmark gene sets and stratified by biological system further confirmed these findings (Supplementary Fig. 2b). Taken together, SIRT6 perturbation produced context-dependent effects across species and biological systems, with mitochondrial dysfunction emerging as the most conserved response to SIRT6 loss.

### 2.3 SIRT6 perturbation shows sex-dependent transcriptional associations with longevity interventions

Next, we decided to evaluate whether SIRT6-associated transcriptional changes recapitulate those observed under established longevity interventions. To investigate this we performed a rank-based GSEA using eight RNA-seq datasets on lifespan- and healthspan-extending interventions in liver from Tyshkovskiy et al. [45]. These signatures were compared with hepatic SIRT6-OE (GSE157838) and SIRT6-KO dataset (GSE129370). Obtained connectivity scores revealed clear sex-specific differences in the SIRT6-OE cohort (Fig. 2d, Supplementary Table 5). In females, all five interventions with available sex-specific gene signatures (Acarbose, 17*α*-estradiol, caloric restriction, Protandim, and Rapamycin) showed negative connectivity, with significant associations for Acarbose, 17*α*-estradiol, and caloric restriction (*p*-value *<* 0.001). In males, the pattern was more variable, with positive connectivity for Acarbose and a weaker positive association with Rapamycin, whereas Protandim and 17*α*-estradiol showed negative connectivity. The SIRT6-KO profile showed an opposing trend relative to female SIRT6-OE for several interventions, reaching significance for Acarbose but not for 17*α*-estradiol or caloric restriction (Supplementary Fig. 2c, Supplementary Table 6). Additionally, analyzed interventions did not significantly affect hepatic SIRT6 expression levels, suggesting that their connectivity with SIRT6-associated changes reflects modulation of downstream transcriptional programs rather than direct induction of SIRT6 (Supplementary Fig. 2d).

### 2.4 SIRT6 perturbation induces divergent TF responses in mouse and human

Given the widespread expression alterations in SIRT6 perturbation models, we next asked which upstream regulators drive these changes and whether they are conserved across species. To address this question, we inferred transcription factor (TF) activity in mouse and human SIRT6 perturbation models. Our analysis revealed reciprocal TF activity patterns in mice: TFs activated upon KO tended to be repressed upon OE (Fig. 2e). Among TFs significant in at least one perturbation condition, 75% showed this mirror response (289 out of 385, binomial *p*-value = 1.4 10*^−^*^23^, Spearman *ρ* = −0.33). This pattern was dominated by regulators of the inflammaging axis, including NF-*κ*B members (RELA, REL, RELB, NFKB1, NFKB2), AP-1 components (JUN, JUNB, JUND, FOS), and STAT1/STAT3.

We next asked whether this TF activity was conserved between mouse and human, and found that it was not: among 66 TFs significant in KO data from both species, only 33% were concordant (binomial *p*-value = 9 10*^−^*^3^). The strongest divergence involved the inflammaging axis (*n* = 13), for which only 2 out of 13 TFs were concordant in their activity in cross-KO comparisons (binomial *p*-value = 0.023), while cross-OE effects showed a Spearman *ρ* = −0.66 (*p*-value = 0.017).

In mouse models, SIRT6 loss consistently activated members of NF-*κ*B, AP-1, and STAT TF families, including JUN (*z_meta_*= 2.16), REL (*z_meta_* = 2.01), NFKB1 (*z_meta_* = 1.86), and STAT3 (*z_meta_* = 1.02) (Supplementary Fig. 3a). In human systems, however, this regulatory axis was repressed, as reflected by reduced activity of STAT3 (*z_meta_* = -1.75), JUND (*z_meta_* = -1.01), and JUN (*z_meta_* = -0.87). JUN, JUND, and STAT3 were significant in both species. Overexpression showed the same split in reverse, repressing NF-*κ*B/AP-1/STAT activity in mouse but increasing it in human cells (Supplementary Fig. 3b). These contrasting responses may reflect differences in baseline NF-*κ*B regulation between *in vivo* mouse tissues and cultured human cell systems.

The remaining aging-related axes (proteostasis/redox, mitochondrial, senescence, nutrient sensing, growth/epigenetic) contained too few TFs (*n* ≤ 6) to support robust species-level inference. Across the knockout data, only two transcription factors were significant and co-directional in both species at the individual level: TP53 (*z_meta_*= 0.67 in mouse, *z_meta_*= 1.32 in human) and ESRRA (*z_meta_* = -0.75 in mouse, *z_meta_*= 1.03 in human) were significant and co-directional in both species, making them the only conserved SIRT6-responsive candidates identified. Notably, these TFs are involved in cellular senescence and mitochondrial dysfunction, the aging hallmarks most extensively described in relation to SIRT6 in our topic modeling analysis (Fig. 1d). MYC, whose transcriptional output has previously been shown to be repressed by SIRT6 [46] also showed significant activity changes in both knockout datasets, but the direction differed between species (*z_meta_*= 0.87 in mouse and *z_meta_*= − 1.06 in human). Together, these findings show that TF responses to SIRT6 perturbations are largely species-specific, with limited conservation beyond TP53 and ESRRA.

### 2.5 Conserved SIRT6-dependent signatures highlight aging-related transcriptional programs

Heterogeneity of gene expression changes across datasets following SIRT6 perturbation motivated us to identify conserved SIRT6-dependent transcriptional signatures across species (Fig. 3a). To identify conserved gene signatures of SIRT6 perturbation, we applied a multivariate meta-regression-based approach. Meta-analysis of SIRT6-KO datasets from six organisms revealed 30 significant genes at FDR-adjusted *p*-value *<* 0.05, including two genes that also showed a log_2_ FC *>* 0.58 (Fig. 3b, Supplementary Fig. 4a, Supplementary Table 7). Interestingly, most of the significant genes showed relatively low heterogeneity (I^2^ *<* 50%), which could indicate consistent direction and magnitude of expression changes across different species and experimental conditions (Fig. 3b). However, low heterogeneity should be carefully interpreted as the value can be underestimated due to the lack of statistical power. A minor subset of significant genes demonstrated high heterogeneity (I^2^ *>* 50%) suggesting that their transcriptional response to SIRT6 deficiency varies depending on biological context, such as tissue or cell type, experimental design and organism. Moreover, 20 of the 30 significant genes retained significance in the leave-one-out analysis (LOO *p*-value *<* 0.05), supporting the robustness of the meta-analysis results (Supplementary Fig. 4b).

**Fig. 3.**
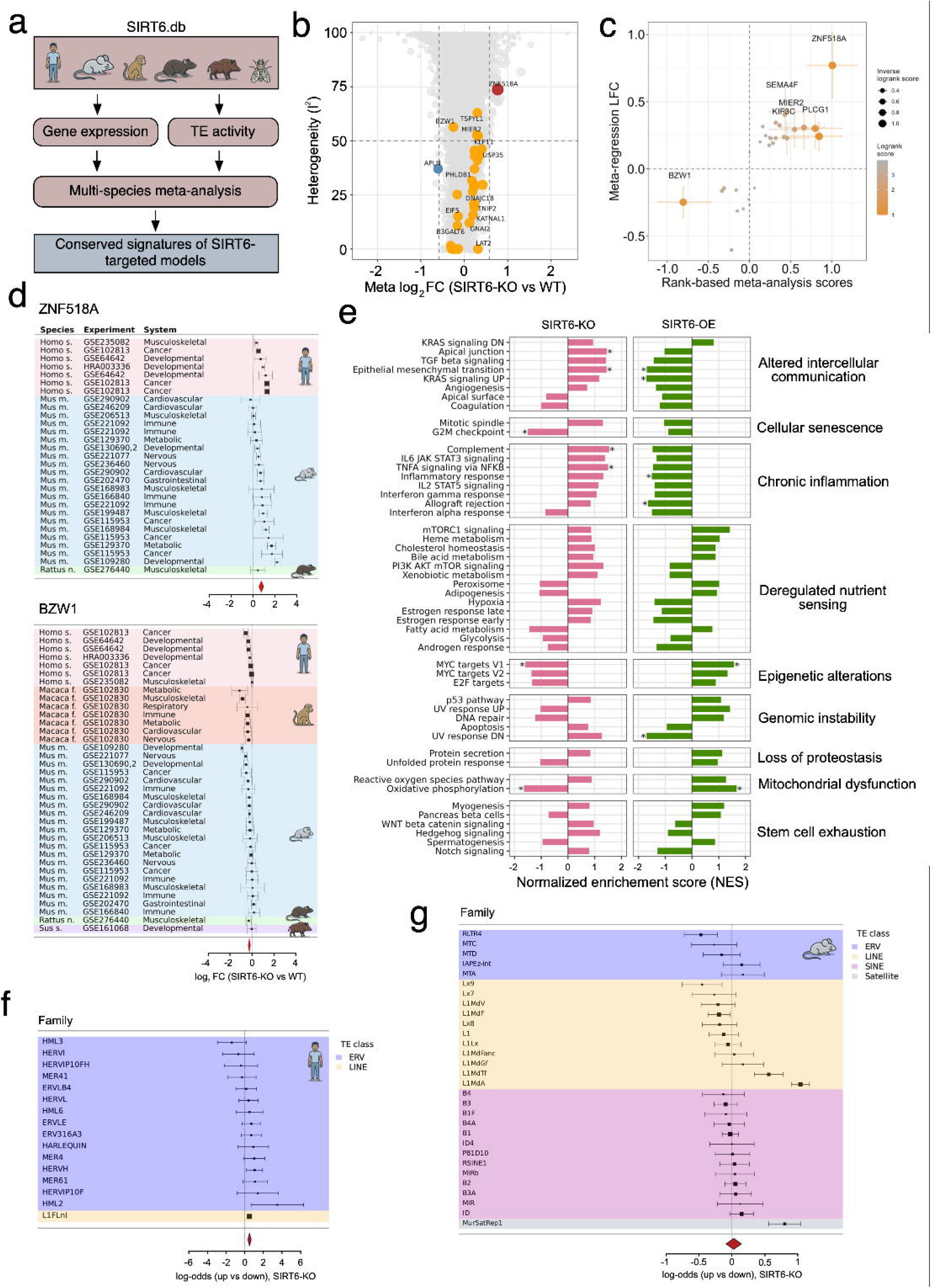
Identification of SIRT6 transcriptional signatures. (a) Overview of the meta-analysis performed using datasets from SIRT6.db. (b) Volcano plot showing the regression-based meta-analysis results for SIRT6-KO datasets across species. Highlighted circles shows genes, statistically significant in a meta-analysis. (c) Scatter plot illustrating the most affected targets identified by two meta-analysis approaches. (d) Forest plots of *ZNF518A* (upper panel) and *BZW1* (lower panel) expression changes across datasets included in the meta-analysis. The size of the squares demonstrates the weight of each individual log_2_FC into meta log_2_FC. The red diamond shows the meta log_2_FC of the gene and the width of the diamond demonstrates the confidence interval (e) Bar plots representing normalized enrichment score (NES) values obtained from GSEA on MSigDB pathways for SIRT6 deficiency and overexpression experiments. Asterisks show FDR *p*-values *<* 0.05. (f-g) Forest plots showing changes in transposable elements family expression in human (left panel) and mouse (right panel).

Given that signatures identified by our meta-regression approach may be affected by differences in the number of species in which a given gene is represented, we sought to validate these results using an independent meta-analysis strategy. For this purpose, we applied a complementary gene-ranking approach inspired by the method of Stephen Gammie [47], which ranks genes by the consistency and strength of their differential expression across datasets and species, prioritizing genes that repeatedly appear among the most strongly up or downregulated (described in Methods). Two genes showed the strongest and most consistent effects across both meta-analysis approaches (Fig. 3c, Supplementary Fig. 4a, Supplementary Table 8). One was upregulated (*ZNF518A*, log_2_FC = 0.77), while the other was downregulated (*BZW1*, log_2_FC = -0.6). Notably, *ZNF518A* was detected in mammals across 3 organisms (human, mouse and rat) with high heterogeneity (I^2^ = 73.7%), indicating that while SIRT6 deficiency consistently upregulated this gene, the magnitude of this effect varies across experimental contexts (Fig. 3d). Transcriptional effects of *ZNF518A* in case of cancer showed the highest and most stable response in the absence of SIRT6, thus, these results made a bigger contribution in identifying meta log_2_FC. The expression of the gene was mostly upregulated in mouse datasets, showing the biggest transcriptional response in the developmental biological system (log_2_FC = 2.22). Interestingly, protein-truncating variants in *ZNF518A* have been linked to shorter female reproductive lifespan [48], although its role in the aging of other biological systems remains to be established. In contrast, despite its modest log_2_FC, *BZW1* was among the most consistently downregulated genes, with downregulation detected in five of the six species examined (Fig. 3d). Functionally, *BZW1* encodes an eIF5-mimic translational regulator of start-codon selection and translation initiation [49], and its dysregulation has been linked primarily to cancer, including tumor growth and metastatic phenotypes [50]. Additionally, using publicly available SIRT6 CUT&RUN data [51], we independently confirmed SIRT6 occupancy at the promoters of the five top-ranked genes (*ZNF518A*, *TSPYL1*, *PLCG1*, *MIER2*, *BZW1* ) prioritized by both meta-analysis approaches, as well as at the *APLN* promoter (Supplementary Fig. 4c).

The meta-analysis of SIRT6-OE vs WT identified 94 significant genes (with FDR-adjusted *p*-value *<* 0.05) and 32 differentially expressed genes among them (Supplementary Fig. 4d). We also detected a predominance of downregulated DEGs (27 genes), which is consistent with the histone deacetylase activity of SIRT6 and its role in transcriptional repression. Most significant genes were detected only in human and mouse datasets, whereas four were also detected in Drosophila but showed small transcriptional effects. Heterogeneity was generally low (*I*^2^ *<* 50%), with moderate heterogeneity observed only for *HIP1* and *KNSTRN* (Supplementary Figure 4d). However, these estimates should be interpreted cautiously given the limited dataset number (10 experiments across three species).

Given the established role of SIRT6 in lifespan regulation and the aging-like phenotype, we decided to compare transcriptional changes identified in SIRT6-KO and SIRT6-OE meta-analyses and map them to the hallmarks of aging. The combined analysis of GSEA results from SIRT6-KO and OE meta-analyses revealed that SIRT6 deficiency and overexpression lead to the opposite transcriptional changes across biological pathways. In total, seven pathways were significant in SIRT6-KO and SIRT6-OE meta-analyses, with three pathways (oxidative phosphorylation, EMT and MYC target V1) reaching significance in both conditions (Fig. 3e). Moreover, oxidative phosphorylation represented the most consistent (NES = -1.64, FDR *p*-value = 2.37 10*^−^*^3^ and NES = 1.67, FDR *p*-value = 1.77 10*^−^*^4^ in SIRT6-KO and SIRT6-OE, respectively) bidirectional pathway across both meta-analyses. A complementary GSEA using Open Genes aging hallmark gene signatures confirmed this bidirectional pattern, with opposite enrichment changes observed for six of the eleven aging hallmarks (Supplementary Fig. 4e). Both these analyses indicate mitochondrial impairment upon SIRT6 deficiency, overall positioning it as a central aging-related process associated with SIRT6 functions.

### 2.6 Transposable elements family expression changes are consistent in humans but heterogeneous in mice

Given the previously published studies on the role of SIRT6 in regulation of transposable elements (TE) in the genome [52, 53], we next sought to identify TE families that are consistently affected by SIRT6 perturbation in the two species represented by the largest number of individual datasets. For each TE family, we summarized the balance of significantly up- versus downregulated loci upon SIRT6 loss and combined these family-level estimates in a meta-analysis (Fig. 3f,g, Supplementary Tables 9,10). In humans (Fig. 3g), nearly all analyzed families (almost exclusively endogenous retroviruses (ERVs)) showed the same clear shift toward upregulation upon SIRT6 loss, with very little variation between families: roughly 62% of significantly changed loci were upregulated across the class as a whole. The most strongly and reliably upregulated families included *HML2* (HERV-K), *L1FLnI*, and *HERVH*. In mice (Fig. 3f), by contrast, there was no overall bias toward up- or downregulation when averaged across all families (close to a 50/50 split). This average obscured substantial differences between individual families, however: some, such as the major satellite repeat *Mur-SatRep1* and the young LINE-1 subfamilies *L1MdA* and *L1MdTf*, were significantly upregulated, while others, including the older LINE-1 subfamily *L1MdF*, *Lx9*, and the ERV family *RLTR4*, were significantly downregulated. Thus, rather than supporting a universal increase in LINE-1 activity upon SIRT6 deficiency, our results indicate that activation across biological systems and experimental models is mostly restricted to selected, evolutionarily younger LINE-1 subfamilies.

### 2.7 SIRT6 expression is generally stable across tissues but declines in fibroblasts in aging

To further assess the potential of SIRT6 as a transcriptomic biomarker of aging, we analyzed multi-tissue human gene expression data in the datasets reported in ARCHS4 database [54] (Fig. 4a). Across most tissues examined, SIRT6 expression showed modest or statistically insignificant age-dependent changes (Fig. 4b, Supplementary Fig. 5a, Supplementary Table 11). Eight comparisons across six datasets were significant at FDR-adjusted *p*-value *<* 0.05: SIRT6 expression was lower in the older group in five contrasts and higher in three. At the same time, SIRT6 was insignificant in all Old vs Young comparisons, arguing against a uniform age-associated change in SIRT6 across tissues.

**Fig. 4.**
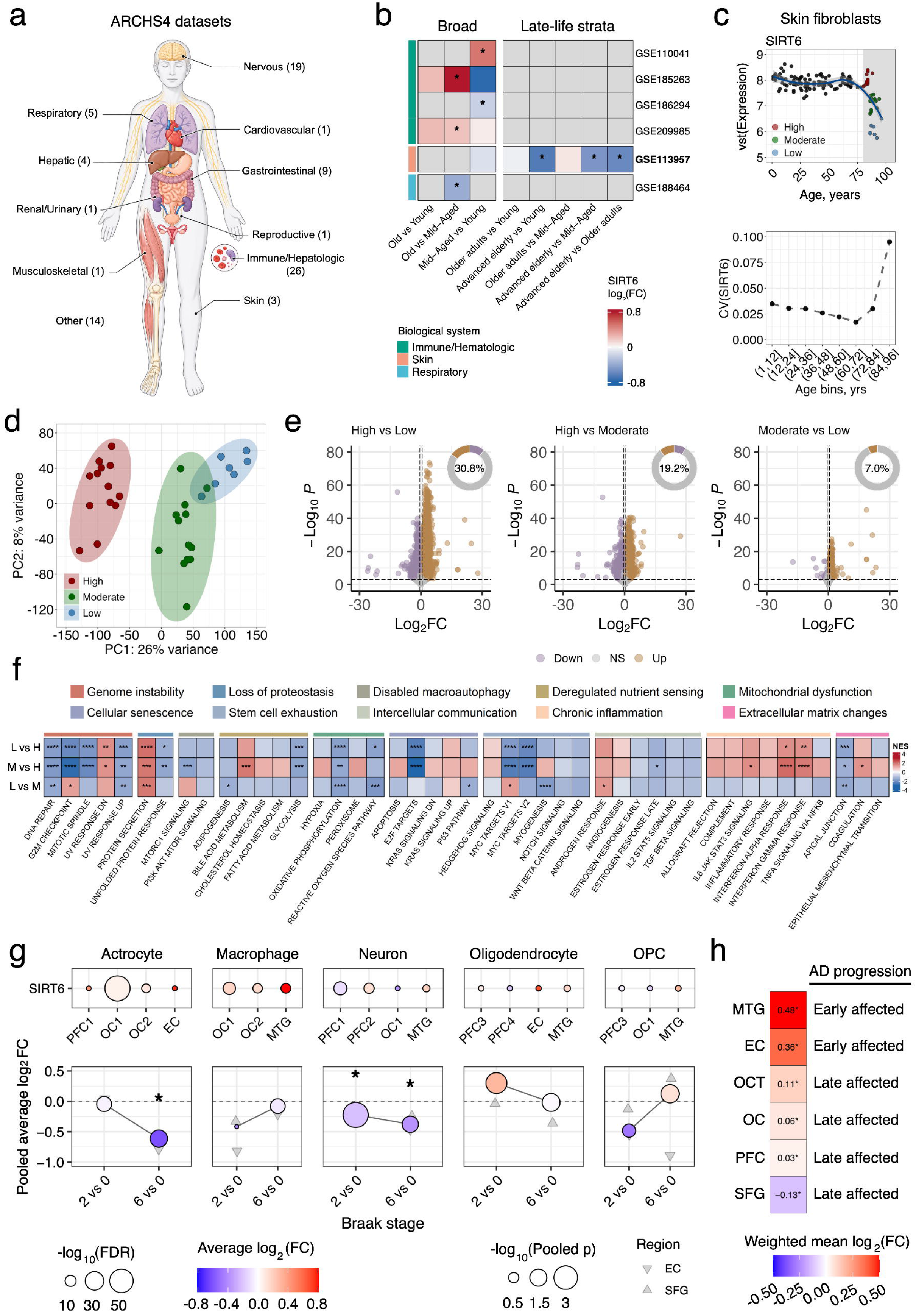
Transcriptional changes in SIRT6 during normal and pathological aging. (a) Biological systems with corresponding number of datasets represented in the selected ARCHS4 experiments with detectable SIRT6 expression. (b) Heatmap of SIRT6 expression changes that were significant in at least one comparison. Asterisks indicate FDR-adjusted *p*-value *<* 0.05. (c) Age-related trajectories of SIRT6 expression (upper panel) and its coefficient of variation (lower panel) in dermal fibroblasts from GSE113957. Red, green, and blue dots represent advanced age donors with high, moderate, and low SIRT6 expression, respectively. (d) PCA of skin fibroblast expression profiles from donors of advanced age, colored by SIRT6 expression level (high, moderate, or low). (e) Volcano plots of differential expression between SIRT6-expression groups (high, moderate, low) in skin fibroblasts. (f) GSEA results for MSigDB Hallmark gene sets across these comparisons, stratified by their relevance to the hallmarks of aging. (g) Cell-type-specific changes in SIRT6 expression across AD datasets. (h) Region-specific changes in SIRT6 expression, represented by the weighted mean log_2_FC.

A notable exception was observed in human skin fibroblasts, where SIRT6 expression was negatively associated with chronological age (partial Spearman *ρ* = −0.377, *p* = 8.29 10*^−^*^6^, Supplementary Fig. 5b), remaining relatively stable through most of adulthood before declining markedly at advanced ages (Supplementary Fig. 5c, Fig. 4c, upper panel). In donors older than 80 years, SIRT6 expression was significantly lower than in young individuals (log_2_FC = −0.672, adjusted *p*-value = 4.19 10*^−^*^4^), middle-aged individuals (log_2_FC = −0.518, adjusted *p*-value = 3.56 10*^−^*^2^), and individuals aged 60-80 years (log_2_FC = −0.591, FDR-adjusted *p*-value = 4.33 10*^−^*^2^). These changes were also accompanied by increased heteroscedasticity in mRNA levels with age (ΔCV = 0.0649), ranking SIRT6 within the top 2.4% of genes showing increased variability in the oldest age range (Supplementary Fig. 5d, Fig. 4c, bottom panel).

Given the pronounced decline in SIRT6 expression in skin fibroblasts from donors older than 80 years, we focused subsequent analyses on this age group to characterize the associated transcriptional changes. Based on SIRT6 expression, these donors were stratified into three groups with high (“High”), moderate (“Moderate”), or low (“Low”) expression levels (Fig. 4c, upper panel). Donors in the High group were younger on average than those in the Moderate and Low groups, consistent with a progressive decline in SIRT6 expression at advanced ages. Principal component analysis of whole-transcriptome profiles independently supported this stratification, with the Moderate and Low groups separating from the High group primarily along PC1 (Fig. 4d). Differential expression analysis identified 6134 genes between the High and Low groups, 4433 genes between the High and Moderate groups, and 1517 genes between the Moderate and Low groups, indicating extensive remodeling of the fibroblast transcriptome associated with reduced SIRT6 expression in advanced age (Fig. 4e, Supplementary Tables 12-14). To map these changes to cellular functions, we further performed GSEA analysis on MSigDB halmark sets (Fig. 4f). The late-life decline in SIRT6 expession in fibroblasts was consistently associated with significant enrichment in protein secretion and enrichment decrease in DNA repair, oxidative phosporylation and apical junction in all three comparisons. Also, in Low vs High and Moderate vs High groups we detected increased NES scores in pathways related to chronic inflammmation, suggesting a consistent activation of inflammatory signaling in aged fibroblasts upon SIRT6 reduction.

Together, these findings indicate that SIRT6 is not a universal transcriptomic biomarker of chronological aging in humans but may reflect a late-life state in skin fibroblasts, characterized by widespread gene-expression changes and dysregulation of key aging-related processes, including DNA repair, oxidative phosphorylation, proteostasis, and inflammatory signaling.

### 2.8 Temporal and regional heterogeneity of SIRT6 expression in Alzheimer’s disease

Multiple pieces of evidence have implicated SIRT6 in the progression of Alzheimer’s disease [55, 56]. To determine whether the SIRT6 changes observed in normal ageing are recapitulated in pathological aging program, we examined SIRT6 expression across ten independent AD scRNA-seq datasets, spanning six cortical regions: prefrontal cortex (PC), superior frontal gyrus (SFG), middle temporal gyrus (MTG), entorhinal cortex (EC), occipital and occipitotemporal cortex (OC and OCT) (Fig. 4g, Supplementary Table 15).

Across AD vs control comparisons, SIRT6 was consistently downregulated in neurons, with the effect detected in the dorsolateral prefrontal cortex (PFC1) and reproduced, though more weakly, in the remaining neuronal datasets. In contrast, SIRT6 was consistently upregulated in microglia/CNS macrophages across all three cohorts in which this population was recovered (OC, OCT, MTG), with the strongest and most significant induction in the MTG. The oligodendrocyte lineage (oligodendrocytes and oligodendrocyte precurson cells) showed no coherent change: effect sizes were small and inconsistent in sign across cohorts. Astrocytes displayed the most heterogeneous behaviour, with effects ranging from essentially null in OC to modest upregulation in EC.

To resolve the temporal ordering of these changes, we exploited the GSE147528 [57], in which SFG and EC were profiled at defined Braak stages, and summarised the two regions per stage using a significance-weighted mean effect and Stouffer-combined significance (Fig. 4g, lower panel). SIRT6 in neurons was significantly reduced already at Braak stage 2 (weighted mean log_2_FC = 0.22, *p*_combined_ *<* 0.05) and remained reduced at Braak stage 6, indicating that neuronal SIRT6 loss is an early event rather than a terminal consequence of neurodegeneration. Astrocytic SIRT6, by contrast, was unchanged at Braak stage 2 and showed a pronounced decrease only at Braak stage 6 (weighted mean log_2_FC = 0.61, *p*_combined_ *<* 0.05), consistent with a stage-dependent, secondary astrocytic response. Microglia, oligodendrocytes and oligodendrocyte precursor cells showed no significant pooled change at either stage, and the individual regional estimates (Fig. 4g) diverged in sign: the glial response is regionally heterogeneous.

Aggregating across the five major cell types, the pooled regional effect on SIRT6 tracked the known anatomical staging of AD (Fig. 4h). The two regions affected earliest in the disease course (MTG, weighted mean log_2_FC = 0.48 and EC, weighted mean log_2_FC = 0.36) showed the strongest SIRT6 upregulation, whereas regions affected later showed progressively weaker effects (with changes of 0.11 for OCT, 0.06 for OC, 0.03 for PFC) or even downregulation (with changes −0.13 for SFG). All regional alterations reached *p*-value_combined_ *<* 0.05. Together, these results show that SIRT6 dysregulation in AD is cell-type and stage-specific, with an early, persistent decline in neurons, a late decline in astrocytes and consistent upregulation in microglia.

## 3 Discussion

In this study, we integrated NLP-based literature analysis with bioinformatic approaches to systematically evaluate SIRT6 as a transcriptomic biomarker of aging. The literature increasingly linked SIRT6 to multiple aging hallmarks, particularly cellular senescence, mitochondrial dysfunction, and telomere attrition, while gene expression analysis indicates that SIRT6 perturbation produced strongly variable transcriptional responses across species, tissues, and experimental systems. Despite this heterogeneity, complementary meta-analysis approaches identified a few conserved gene signatures, including the upregulation of *ZNF518A* and downregulation of *BZW1*, and revealed reciprocal effects of SIRT6 loss and overexpression on oxidative phosphorylation and inflammatory signaling (Fig. 5). SIRT6 deficiency also produced species-specific effect on TE activity, with broad ERV derepression in human datasets but heterogeneous family-level effects in mice. Finally, SIRT6 expression was largely stable during normal human aging, whereas Alzheimer’s disease datasets showed significant, but cell-type, region and stage-specific dysregulation.

**Fig. 5.**
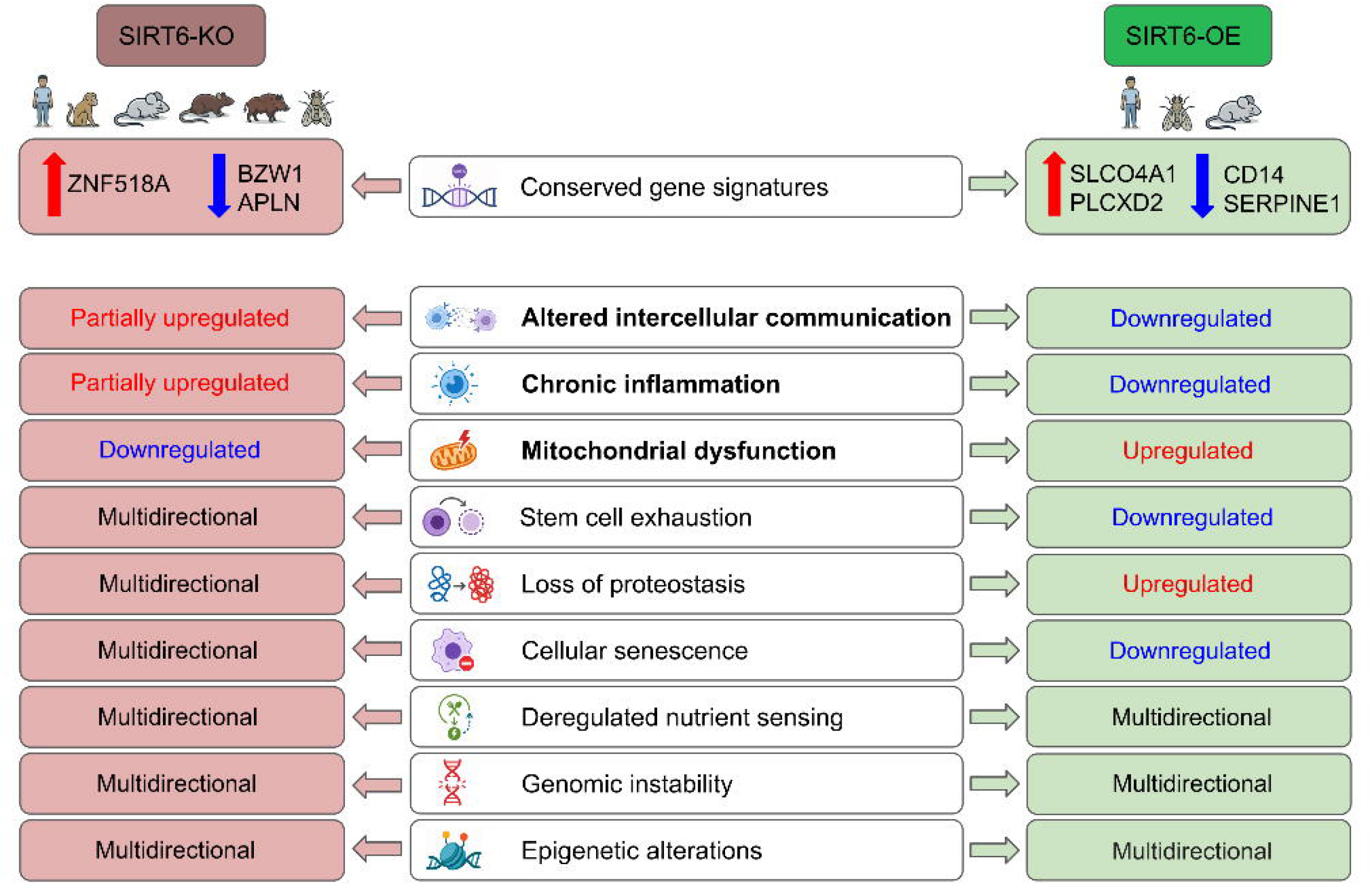
Summary of SIRT6-associated transcriptomic signatures and its relationship with the hallmarks of aging.

A major caveat in interpreting results from bulk transcriptomic data is that each sample represents a mixture of cell populations whose relative abundance may change with age, disease, or experimental perturbation. Consequently, an apparent change in SIRT6 expression or in a SIRT6-associated pathway may reflect altered cell-type composition rather than a cell-intrinsic transcriptional response. This issue is particularly relevant in aging tissues, where immune infiltration, fibrosis, neuronal loss and changes in stromal populations can substantially reshape tissue composition. Our single-cell analysis of AD donors clearly illustrates this problem: SIRT6 was generally downregulated in neurons, but upregulated in microglia/CNS macrophages, meaning that these opposing cell-type-specific effects could be attenuated or even reversed in bulk tissue measurements depending on the cellular composition of each sample.

A second limitation arises from biological differences among the genetic models grouped under the broad labels of SIRT6 knockout or overexpression. The analyzed experiments included models that differed in perturbation strategy, developmental timing, tissue specificity, genetic background, and the duration of SIRT6 depletion or overexpression. Moreover, nominally equivalent knockout models may retain different amounts of SIRT6 RNA or protein, and partial depletion may produce substantially weaker effects than complete loss. Consistent with this interpretation, the proportion of differentially expressed genes varied widely among datasets, with relatively modest responses in many mouse experiments but extensive transcriptional reprogramming in SIRT6-deficient developmental cells. Thus, comparisons across models should account for the measured magnitude of SIRT6 perturbation, rather than treating knockout or overexpression status as a uniform binary condition.

SIRT6-associated transcriptional responses were also strongly tissue dependent. In *Macaca fascicularis*, profiling the same SIRT6 knockout across six tissues revealed that mitochondrial dysfunction and disabled macroautophagy were enriched only in selected tissues, whereas epigenetic alterations were detected in a different subset. Similarly, SIRT6 expression remained largely stable across most tissues during normal human aging, with a notable decrease observed in skin fibroblasts from donors older than 80 years. Regional analyses of AD donor data further showed strong SIRT6 upregulation in the MTG and EC but weak or negative effects in later-affected regions, including downregulation in the superior frontal gyrus. These findings indicate that neither the magnitude nor the direction of a SIRT6-associated signal can be assumed to generalize across tissues or anatomical regions.

Finally, differences between cultured cell models and postmortem tissues complicate direct comparisons across datasets. Cell lines and primary cultures provide relatively homogeneous and experimentally controlled systems, but culture conditions alter proliferation, metabolism, stress responses, and cell-cell interactions, potentially modifying the transcriptional consequences of SIRT6 perturbation. Postmortem tissues better preserve disease-associated cellular environments, yet they introduce additional variation related to cell-type composition, agonal state, postmortem interval, RNA integrity, and clinical heterogeneity. In our analysis, NF-*κ*B-, AP-1-, and STAT-related transcription factor activities showed opposite responses between predominantly *in vivo* mouse tissues and cultured human systems, whereas postmortem AD datasets revealed marked cell-type and region-specific SIRT6 dysregulation. Because species, experimental setting, and sample source are partly confounded in the available data, such differences should not be attributed solely to evolutionary divergence without validation in matched biological systems.

## 4 Methods

### 4.1 Abstract corpus collection

SIRT6-related publications were retrieved from OpenAlex database using PyAlex package by searching article titles and abstracts published since 1950 for the “SIRT6” “Sirt6” “sirt6” “Sirtuin 6” and “Sirtuin6” keywords. Results from the title and abstract searches were combined, retracted records were excluded. Only articles and preprints were retained in the output and records with duplicate titles were removed, yielding 2025 publications. PubMed identifiers were standardized and, when missing, recovered from DOIs using the NCBI Entrez ESearch API. Very short abstracts containing fewer than 20 words or fewer than two sentences were considered incomplete and were re-retrieved from PubMed using EFetch. Anomalously short titles were recovered in the same manner. Additionally, records with titles indicative of withdrawn, supplementary, figure, or review-only material were excluded. Abstract text was subsequently normalized by removing duplicated titles, URLs, email addresses, citations, publication metadata, disclosure and reference sections, trial-registration information, and other non-native patterns for abstracts. Unicode, HTML entities, spacing, and SIRT/NAD nomenclature were standardized, and sentence-level language detection was used to retain English-language text. Finally, extremely similar abstracts were identified using TF–IDF representations with English stop words and removed when pairwise cosine similarity exceeded 0.95, resulting in a final corpus of 1648 abstracts.

### 4.2 SIRT6 centrality analysis

To measure whether SIRT6 is a focus or inciderntal element of each publication, we calculated a title and abstract-based centrality scores ranging from 0 (incidental) to 1 (primary focus). The SIRT6 centrality score, accounting for title presence, mention density, sentence focus, functional context and abstract position was calculated as

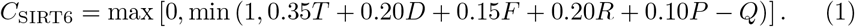

where *T* was 1 when a SIRT6 variant occurred in the title and 0 otherwise *D* = min(1, 40*M/W* ), where *M* is the total number of mentions and *W* is the combined title and abstract word count; and *F* = min[1, (*N*_SIRT6_*/N*_all_)*/*0.55], where *N*_SIRT6_ and *N*_all_ are the numbers of SIRT6-containing and total abstract sentences. Functional context was defined as *R* = *N*_role_*/N*_SIRT6_ when *N*_SIRT6_ *>* 0, and 0 otherwise. Abstract position was defined as *P* = (*I*_first_ + *I*_last_)*/*2, with each indicator equal to 1 when the corresponding sentence contained SIRT6.

The incidental-mention penalty was

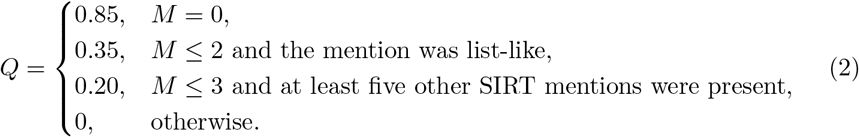

The weighted centrality score was clipped to the interval [0, 1]. Scores were calculated for all papers using the same rules.

### 4.3 Topic modeling analysis

For topic modeling, a corpus comprising 1,648 abstracts of SIRT6-related scientific publications was compiled from the PubMed and OpenAlex databases based on the presence of SIRT6 mentions in the text. Topic modeling, both in the standard and zero-shot settings, was performed using the BERTopic framework [58] with the embeddinggemma-300M model to generate vector representations of the concatenated titles and abstracts. Embeddings were reduced to five dimensions using UMAP, followed by HDBSCAN clustering with a minimum cluster size of 25. Topic terms were generated using a count vectorizer containing unigrams and bigrams occurring in at least two documents. Standard English stop words and frequently occurring domain terms, such as “SIRT6”, “sirtuin”, “activity”, “expression”, “protein”, and “gene”, were excluded. Topic representations were calculated using class-based TF–IDF with BM25 weighting and frequent-word reduction. Next topics were refined using maximal marginal relevance method with a diversity parameter of 0.3.

In a zero-shot analysis, abstracts were mapped to 14 predefined aging hallmarks represented by expanded semantic descriptions, using a minimum cosine-similarity threshold of 0.3. Unmatched documents were clustered with HDBSCAN using a minimum cluster size of 10 documents. Topic probabilities and annual topic frequencies were subsequently calculated from publication years. Finally, a hallmark fingerprint was generated for each abstract by computing cosine similarities between normalized abstract and hallmark-description embeddings.

### 4.4 Collection of SIRT6-targeted RNA-seq datasets

Raw bulk RNA-sequencing data from 28 datasets comprising *Homo sapiens*, *Mus musculus*, *Rattus norvegicus*, *Macaca fascicularis*, *Sus scrofa*, and *Drosophila melanogaster* experiments were retrieved from the GEO National Center for Biotechnology Information (NCBI) using the nf-core/fetchngs v1.12.0 pipeline (https://doi.org/10.5281/zenodo.10728509). The last dataset in the collection (HRA003336) was downloaded from Genome Sequence Archive of the National Genomics Data Center, China.

### 4.5 RNA-seq data preprocessing

RNA-seq data were processed using the nf-core/rnaseq v3.21.0 pipeline (https://doi.org/10.5281/zenodo.20072251). FASTQ quality was assessed with FastQC, followed by adapter trimming and quality filtering (Phred score *<*20) with Trim Galore. Reads were mapped to the corresponding reference genome, and gene-level abundance was quantified using Salmon pseudo-alignment. Reference genomes were GRCm39 for *Mus musculus*, hg38 for *Homo sapiens*, GRCr8 for *Rattus norvegicus*, ensemble genome Macaca fascicularis for *Macaca fascicularis*, BDGP6.54 for *Drosophila melanogaster*, and ensemble genome Sscrofa11 for *Sus scrofa*.

### 4.6 Differential gene expression analysis

Differential expression analysis was performed using an R script made available on GitHub (https://github.com/SIRT6/Reconsidering-the-role-of-sirt6-in-aging/tree/main/SIRT6_DE). Lowly expressed genes were removed by retaining genes with at least 10 raw counts in at least two samples or half of the samples in each group, whichever was greater. Tissues and cell types were manually grouped into nine broader biological systems.

In DE analysis, experiments were stratified by treatment, cell type, tissue, or condition when applicable and incorporated relevant biological covariates, including sex, age, and strain, into the model formula. Differentially expressed genes were defined using two criteria: FDR-adjusted *p*-value *<* 0.05 and | log_2_ FC| *>* 0.58.

### 4.7 Comparative analysis of SIRT6 perturbation and longevity signatures

Hepatic SIRT6-KO and SIRT6-OE profiles were compared with longevity-intervention signatures from Tyshkovskiy et al. using GSEA-based connectivity analysis [45, 59]. Longevity intervention signatures from the liver of 5-7-month old mice were selected to match hepatic SIRT6-OE samples (GSE157838) and SIRT6-KO dataset (GSE129370, female samples only).

For each intervention, differentially expressed genes from the corresponding DESeq2 contrast versus age and sex-matched controls were separated into up- and downregulated sets using a significance threshold of adjusted *p*-value *<* 0.05. When fewer than 15 genes reached significance, the top 150 genes ranked by nominal *p*-value were used. SIRT6 perturbation genes (SIRT6-OE vs. WT, analyzed separately by sex; SIRT6-KO vs. WT, females) were ranked using the signed statistic:

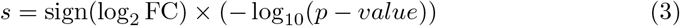

and intervention gene sets were tested using fgsea [60].

NES was used to quantify the strength and direction of gene-set enrichment within each ranked SIRT6 signature. Connectivity was calculated as the average of the NES for the upregulated set and the sign-inverted NES for the downregulated set:

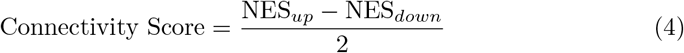

Positive scores indicated concordant transcriptional responses between the SIRT6 perturbation and the corresponding longevity intervention, whereas negative scores indicated opposing responses. Combined significance was calculated by converting the one-sided GSEA *p*-values into signed *z* -scores and combining them as 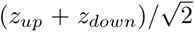. The corresponding *p*-value was obtained from the standard normal distribution. For two comparisons (GHRKO and Snell dwarf signatures against the male SIRT6-OE profile), fgsea was not able compute NES and *p*-values with the requested precision (eps = 0) due to pronounced imbalance between positive and negative gene-level statistics in the target ranked profile. These comparisons are reported as NA in Supplementary Table 5 and excluded from the corresponding Supplementary Fig. 2c.

Sirtuin family expression (*Sirt1-Sirt7* ) was examined directly across the same longevity-intervention datasets: for each gene and intervention, log_2_FC estimates from age- and sex-stratified DESeq2 contrasts were pooled using a random-effects meta-analysis (metafor, REML estimator of between-stratum variance *τ* ^2^) to obtain a single aggregated effect size and standard error per intervention [61].

### 4.8 Analysis of TFs regulating SIRT6 targets

Transcription factor activity was inferred using the univariate linear model implemented in the decoupleR R package [62], with the CollecTRI regulatory network as prior knowledge. Each interaction was assigned a mode of regulation of +1 for stimulatory and 1 for inhibitory edges. Ambiguous or duplicate interactions were discarded. Input for each dataset consisted of signed Wald statistics (*stat*) from DESeq2 [63] differential-expression contrasts (SIRT6-KO or OE vs. WT), yielding a single activity score per TF and eliminating.

Per-dataset TF scores were combined within each species using Stouffer’s method: two-sided *p*-values were converted to signed *z*-scores (*z* = sign(score) · Φ*^−^*^1^(1 − *p/*2)) and aggregated as 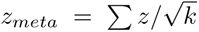, by Benjamini–Hochberg correction. Directional agreement between conditions and species was quantified via binomial sign tests and Spearman’s rank correlation, with analyses restricted to TFs significant (Benjamini-Hochberg adjusted *p*-value *<* 0.05) in at least one or both comparisons.

### 4.9 Regresssion-based meta-analysis of transcriptomic changes in SIRT6-targeted experiments

Firstly, the table with orthologs was prepared for coherent mapping between *Homo sapiens* genes and genes from other 5 model organisms (*Mus musculus*, *Rattus norvegicus*, *Drosophila melanogaster*, *Macaca fascicularis*, *Sus scrofa*). The latest Ensembl version 115 was used to construct an ortholog table by biomaRt R package [64, 65]. Genes with no orthologs were removed, only protein-coding (99.53% of all genes) 1:1 orthologs were kept. There were 18340 genes, 54 experiments and 6 organisms in the final table for meta-analysis.

Secondly, a random-effects multi-level meta-analysis model was applied to find patterns in gene expression changes across the model organisms. The analysis was performed by using rma.mv() function from metafor R package [61]. Dataset ID, species, and phylogenetic relationship between the organisms were used in the model as random terms to extract robust aging signatures, while controlling for batch effects and unequal representation of tissue/species. The phylogenetic correlation matrix was constructed from a phylogenetic tree based on particular divergence time of the organisms (https://timetree.org/). Dataset ID and species random factors were used in nested format to take into account the variance between species and the variance between experiments within species.

Meta-analysis was performed across SIRT6-KO vs WT and SIRT6-OE/K3R OE vs WT experiments separately to identify multi-species gene signatures. For KO meta-analysis, genes that are present in at least 3 organisms and 2 experiments were kept in the input table. Overall, 13359 unique genes from 39 experiments and 6 organisms (*Homo sapiens*, *Mus musculus*, *Rattus norvegicus*, *Macaca fascicularis*, *Sus scrofa*, and *Drosophila melanogaster* ) were tested in a random-effects model. Due to the data availability, genes that are present in at least 2 organisms and 2 experiments were kept in the input table for OE meta-analysis. As a result, 9916 unique genes from 10 experiments and 3 organisms (*Homo sapiens*, *Mus musculus*, and *Drosophila melanogaster* ) were tested in a random-effects model.

### 4.10 Gene set enrichment analysis of aging hallmarks

GSEA on significant genes from individual datasets was performed using the fgsea R package with the fgseaMultilevel [66] algorithm. Aging hallmark gene sets were derived from the Open Genes database [44]. Genes annotated with multiple hallmarks were assigned to each corresponding set. Human HGNC symbols were mapped to species-specific gene symbols using a curated one-to-one ortholog table, retaining only one-to-one relationships and protein-coding genes. The original Open Genes aging mechanism categories were consolidated into the canonical hallmarks of aging framework. Specifically, “degradation of proteolytic systems” and “impairment of protein folding and stability” were merged into *Loss of proteostasis*; “impairment of mitochondrial integrity and biogenesis” and “accumulation of reactive oxygen species” into *Mitochondrial dysfunction*; “chromatin remodeling”, “alterations in histone modifications”, and “transcriptional alterations” into *Epigenetic alterations*; “sterile inflammation” into *Chronic inflammation*; “nuclear DNA instability” into *Genome instability* ; “senescent cells accumulation” into *Cellular senescence*; “changes in the extracellular matrix structure” into *Extracellular matrix changes*; and “intercellular communication impairment” into *Intercellular communication*. For each dataset, genes were filtered to those passing significance thresholds (adjusted *p*-value *<* 0.05 and log_2_ FC *>* 0.58). Pathways with Benjamini–Hochberg adjusted *p*-value *<* 0.05 were considered significantly enriched. All datasets were mapped to a common human gene space and analysed against the human CollecTRI network [67]. Mouse Ensembl identifiers were mapped to human gene symbols via a one-to-one orthologue table. Human identifiers were mapped using org.Hs.eg.db [68].

GSEA on meta-analysis gene signatures was conducted using gene sets from the MSigDB, including Hallmark gene sets. Genes were ranked using a combined metric as in Eq. (3). Pathways were considered enriched based on NES and FDR-adjusted *p*-values. Positive NES values indicated pathways enriched among upregulated genes, while negative NES values corresponded to pathways enriched among downregulated genes. All GSEA hallmarks were mapped to the hallmarks of aging using Claude Opus 4.6 extended AI model, with significant pathways marked for each meta-analysis.

### 4.11 Cross-species ortholog mapping

Pairwise ortholog relationships among all six species were retrieved using the gorth() function from the gprofiler2 R package [69]. We retained only genes present in the SIRT6.db differential expression datasets and restricted the analysis to one-to-one orthologs. Gene IDs were standardized using the following priority order: *Homo sapiens Mus musculus Macaca fascicularis Rattus norvegicus Sus scrofa Drosophila melanogaster*. Genes were assigned the Ensembl identifier of their human ortholog when available. Otherwise, the identifier from the highest-priority available species was used.

### 4.12 Rank-based meta-analysis of cross-species transcriptomic changes

Gene ranking was adapted from the gene expression portrait framework proposed by Stephen C. Gammie [47]. The original approach was extended to integrate transcriptomic datasets from multiple species by incorporating one-to-one ortholog mapping, phylogenetically informed species weighting and bootstrap estimation of confidence intervals.

Within each dataset, differential expression results were first converted into a signed score per gene, as in Eq. (3). where adjusted *p*-values equal to zero were replaced with the smallest non-zero adjusted *p*-value observed in that dataset.

For each organism, genes were ranked independently in upregulated and downregulated directions. Following the original method, the numbers of appearances within the top 1000-8000 ranked genes (step size 1000) were counted and combined using exponentially decreasing weights (1, 0.1, 0.01*, . . . ,* 10*^−^*^7^). The resulting score was normalized by the theoretical maximum possible score and multiplied by the square root of the number of datasets. This normalization prevented overrepresented species, such as mouse, from dominating the final ranking.

Before integrating per-organism scores across all species, orthologs from non-human species were mapped to the unified identifier space using procedure described above. Species contributions were weighted by both phylogenetic distinctiveness and dataset availability.

Phylogenetic weights were calculated using the fair proportion method (evol.distinct(), picante R package [70]) on a dated species tree. Let *f_i_* denote the fair proportion value for species *i*. To reduce overrepresentation of phylogenetically isolated species with limited data, fair proportion values were square-root transformed:

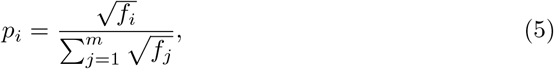

where *m* is the number of species included in the analysis. Dataset availability weights were defined as:

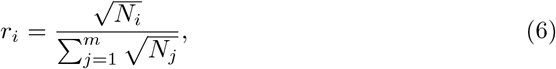

where *N_i_* is the number of datasets available for species *i*. The final combined weight for each species was:

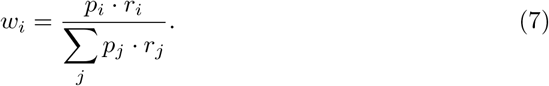

The final conservation score was computed as this weighted sum, with its sign indicating consistent upregulation (positive) or downregulation (negative) across SIRT6-KO models.

Confidence intervals were estimated by bootstrap resampling of datasets within each species (100 iterations). In each iteration, datasets were sampled with replacement, organism-specific scores and the final conservation score were recalculated. 95% confidence intervals were obtained from the obtained bootstrap distribution.

For organism-specific analyses (*Homo sapiens*, *Mus musculus*, and *Macaca fascicularis*), the same ranking procedure was applied independently to datasets from each species. In these analyses, ortholog mapping and cross-species weighting were omitted, and confidence intervals were estimated by bootstrap resampling of datasets within each species (100 iterations), analogous to the cross-species analysis.

### 4.13 Analysis of public CUT&RUN data

Raw CUT&RUN sequencing data on SIRT6 binding sites were obtained from the GSA repository (accession HRA005392) and processed using the nf-core/cutandrun v3.2.2 pipeline (https://doi.org/10.5281/zenodo.5653535) to generate signal tracks. Briefly, raw reads were trimmed, aligned to the hg38 reference genome using Bowtie2 [71], QC filtered and CPM normalized. Signal profiles for selected genes were visualized using the plotgardener R package [72].

### 4.14 Analysis sirtuin expression trajectories in human aging

Human gene-expression counts and sample metadata were obtained from the ARCHS4 human gene v2.7 HDF5 resource. Analyses were restricted to samples explicitly annotated as human and healthy, normal, unaffected, disease-free or control. Samples with missing, ambiguous or disease annotations were excluded. Ages reported in years, months, weeks or days were converted to years. Samples with missing information about donor’s chronological age were discarded. Single-cell and single-nucleus RNA-seq studies were excluded from the analysis. Datasets with missing or duplicated sample identifiers were removed. Samples were stratified to young (*<*30 years), mid-aged (30–60 years) or old (*>*60 years) age groups. Where supported by the data, the old group was subdivided into older adults (*>*60–80 years), advanced elderly (*>*80–100 years) and individuals older than 100 years. A dataset was retained only if at least two age groups each contained three or more healthy samples. Groups not satysfying this threshold were removed before DE analysis.

Differential expression was evaluated separately within each dataset and for every eligible pair of age groups using DESeq2. Genes were retained for a contrast when they had at least 10 counts in at least the greater of two samples or half of the samples in each comparison group. DESeq2 models included age group and, when available sex, ethnicity. Benjamini–Hochberg adjusted *p*-values *<* 0.05 were considered significant. For the DE analysis of GSE113957 human fibroblast dataset ten HGPS samples and genes with ≤ 100 total reads across the retained samples were excluded and inconsistent metadata labels were harmonized. Associations between individual sirtuin expression levels and age were further assessed using partial Spearman correlation, with sex included as a covariate.

To quantify age-associated changes in expression variability, samples were divided into 12-year age intervals. Within each interval, the coefficient of variation was calculated as the standard deviation divided by the absolute mean of variance-stabilized expression. For each gene, the late-life change in variability (ΔCV) was defined as the difference between the coefficients of variation in the 84–96 and 72–84 year intervals. Samples from donors aged ≥ 80 years were selected for analysis of late-life SIRT6 heterogeneity. These samples were divided into groups named ‘High’, ‘Moderate’, and ‘Low’ based on SIRT6 level by Ckmeans.1d.dp [73] clustering method. Differential expression among the these groups of donors was analyzed using DESeq2 with the *design sex* + *age* + *SIRT* 6 *group*, age values were scaled before model fitting. Genes were classified as differentially expressed at adjusted *p <* 0.001 and an absolute shrunken log_2_ FC *>* 0.58. GSEA was performed for the Low vs High, Medium vs High, and Low vs Medium comparisons using the 50 human MSigDB Hallmark gene sets obtained with the msigdbr R package.

### 4.15 Analysis of transcriptional bias by gene length

To test whether SIRT6 loss recapitulates the gene length-dependent transcription decline (GLTD) characteristic of progeroid and aging models, a cross-species meta-analysis of publicly available SIRT6-KO transcriptomic datasets spanning six species (*Homo sapiens*, *Mus musculus*, *Rattus norvegicus*, *Sus scrofa*, *Drosophila melanogaster*, *Macaca fascicularis*) was performed. For each dataset, differential expression results (DESeq2 [63] stat, log2FoldChange, lfcSE, baseMean, padj) were intersected with protein-coding gene models (Ensembl release 110 [74]; autosomal genes only, genomic span ≥ 1 kb) to obtain gene length (log_10_-transformed). Genes with baseMean *<* 20 were excluded to avoid confounding between gene length and expression level. For each contrast, a weighted linear regression

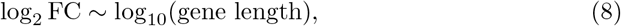

was fitted with weights 1*/*lfcSE^2^, yielding a per-dataset GLTD slope and its standard error (negative slope: preferential downregulation of long genes). A secondary model additionally adjusted for log_10_(baseMean) to control for residual length-expression confounding. Datasets were restricted to SIRT6-KO vs. WT contrasts. Two mouse datasets (GSE206513, GSE221077) were excluded *a priori* as technical artefacts (near-unity *padj* or absence of significant differential expression), independent of GLTD results. For multi-condition datasets (GSE290902, GSE129370, GSE221092 in mouse; GSE102813 in human; GSE309387 in *Drosophila*), only samples without treatment were retained. Per-dataset slopes were combined within each species using random-effects meta-analysis (REML, metafor [61]). Pooling was performed only for species represented by ≥ 2 independent experiments (i.e., distinct GEO/HRA accessions, not distinct tissues or strata within one accession) to avoid pseudoreplication. An overall cross-species pooled estimate was computed analogously. Heterogeneity was quantified using *I*^2^ and *τ* ^2^.

### 4.16 Transposable element quantification and differential expression

Locus-specific transposable element expression was quantified with Telescope [75] using telescope assign, which reassigns multi-mapping reads to individual TE loci via an EM-based Bayesian mixture model. Per-sample count values were merged into a locus-by-sample count matrix (missing loci set to zero, values rounded to integers). Sample metadata and sample-to-experiment mappings were taken from the curated SIRT6 database, and *Homo sapiens* and *Mus musculus* were analysed separately. Differential expression was assessed independently within each experiment using DESeq2 [63] to avoid cross-study batch effects. Experiments were retained only if a WT condition was present, with≥ 2 genotypes, ≥ 2 replicates per genotype, and ≥ 4 samples total. Cell type was added as a covariate (∼ cell type + genotype) when balanced, otherwise the design was ~ genotype. Non-WT genotypes were contrasted against WT with lfcShrink (type = “normal”) and loci with FDR-adjusted *p*-value *<* 0.05 were considered significant. TE loci were assigned to a family and a class (ERV, LINE, SINE, SVA/Satellite, DNA, Other). For the SIRT6-KO contrast, significant loci were aggregated by family, and the proportion of upregulated loci was modelled as a logit effect size (metafor::escalc [61], measure = “PLO”, with Haldane-Anscombe correction). Families with ≥ 5 significant loci were combined in a random-effects meta-analysis (metafor::rma [61], REML). Pooled directional bias was reported alongside Cochran’s *Q*, *I*^2^, and *τ* ^2^.

### 4.17 Cross-cohort analysis of SIRT6 expression in Alzheimer’s disease

#### 4.17.1 Datasets and cell type harmonisation

Publicly available single-cell and single-nucleus RNA-sequencing datasets of postmortem human cortex from Alzheimer’s disease patients and non-demented controls were obtained from GSE214979 [76, 77] (dorsolateral prefrontal cortex), GSE174367 [78] (prefrontal cortex), GSE157827 [79, 80] (prefrontal cortex), GSE129308 [81, 82] (prefrontal cortex, BA9), GSE129308 [83] (occipital and occipitotemporal cortex), GSE138852 [84] (entorhinal cortex), GSE188545 [85] (middle temporal gyrus), and GSE147528 [57] (superior frontal gyrus and entorhinal cortex, stratified by Braak stage). Cell type labels were assigned independently in each study using different marker panels and clustering resolutions; all datasets were therefore re-annotated against a single common reference to make cell type identities directly comparable across cohorts. The Siletti *et al.* (2023) [86] cortical atlas was used as the reference. Both the reference and each query dataset were log-normalised (scuttle::logNormCounts) [87], and cell type labels were transferred using SingleR [88] the reference cell type annotation as the label set. All downstream analyses were performed on these harmonised labels.

#### 4.17.2 Quality control and differential expression

For each dataset independently, genes with zero counts across all cells were removed. Per-cell quality control metrics were computed with scater::perCellQCMetrics [87], and cells whose number of detected genes deviated by more than two median absolute deviations from the median (on the log scale) were excluded. Lowly expressed genes, defined as those detected with more than one count in fewer than ten cells, were removed. Cell types represented by a single cell in any condition were excluded to avoid degenerate comparisons. Differential expression between AD and control cells was computed separately within each cell type using the Wilcoxon rank-sum test as implemented in the Libra framework [89] (run de with de family = “singlecell”, de method = “wilcox”), with sample identity supplied as the replicate variable. For the GSE147528 dataset, comparisons were performed separately for Braak stage 2 versus 0 and Braak stage 6 versus 0 within each of the two profiled regions. *p*-values were corrected for multiple testing across genes, and the resulting per-cell-type tables of log_2_FC (avg logFC) and adjusted *p*-values (p val adj) were retained for SIRT6. Minor cell populations (pericytes, fibroblasts, endothelial cells, and leukocytes) were excluded from all downstream summaries due to their small cell numbers and unstable effect size estimates.

#### 4.17.3 Cross-cohort summarisation

Since the Wilcoxon rank-sum test does not return a standard error for the log_2_FC, a formal inverse-variance meta-analysis was not performed. Two descriptive summaries were used instead.

##### Pooled effect size

For each grouping (cell type × Braak stage for the GSE147528 dataset’s progression analysis; brain region for the regional summary), the pooled effect was computed as a significance-weighted mean of the observed fold changes across the contributing experiments:

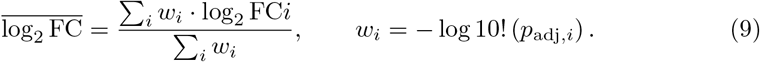

Weights reflect statistical confidence rather than inverse variance.

##### Combined significance

Adjusted *p*-values were converted to signed *z*-statistics,

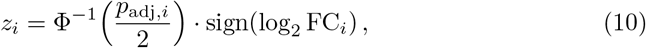

and combined across experiments using the weighted Stouffer method with the same weights *w_i_*:

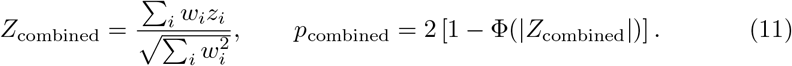

*p*-values were bounded away from 0 and 1 prior to transformation to avoid infinite *z*-statistics. For the GSE147528 dataset’s progression analysis, pooling was restricted to cell type Braak stage combinations for which both regions (SFG and EC) were represented (*k* = 2).

### 4.18 Use of AI-assisted technologies

During the preparation of this manuscript, ChatGPT and Claude were used for language editing, readability improvement, and code review. All AI-assisted suggestions were reviewed and verified by the authors, who take full responsibility for the final content.

### 4.19 Reporting summary

Further information on research design is available in the Nature Portfolio Reporting Summary linked to this article.

## 5 Data availability

The complete set of bulk RNA-sequencing datasets on SIRT6-perturbations used in the study is summarized in Supplementary Table 16. Additionally, ten previously published scRNA-seq datasets were utilized for the analysis of SIRT6 acitivy in AD are summarized in Supplementary Table 15.

## 6 Code availability

The code used for the analyses in this study is available on GitHub: https://github.com/SIRT6/Reconsidering-the-role-of-sirt6-in-aging. The SIRT6.db web app, which can be used to access the processed data and main results in this study is accessible at https://sirt6.github.io/SIRT6.db/.

## 8 Author contributions

E.Ka., A.T, A.M, performed bioinformatic analysis and wrote the paper, N.K. performed bioinformatic analysis, A.P. peformed processing of RNA-seq data, D.T. guided the analysis, D.S conceptualized and planned the project, performed bioinformatic analysis and wrote the paper, E.Kh. guided bioinformatic analysis, wrote the paper and planned the project.

## 10 Additional information

