## Supplementary Figures for "A systematic evaluation of SIRT6 as a transcriptomic biomarker of aging"

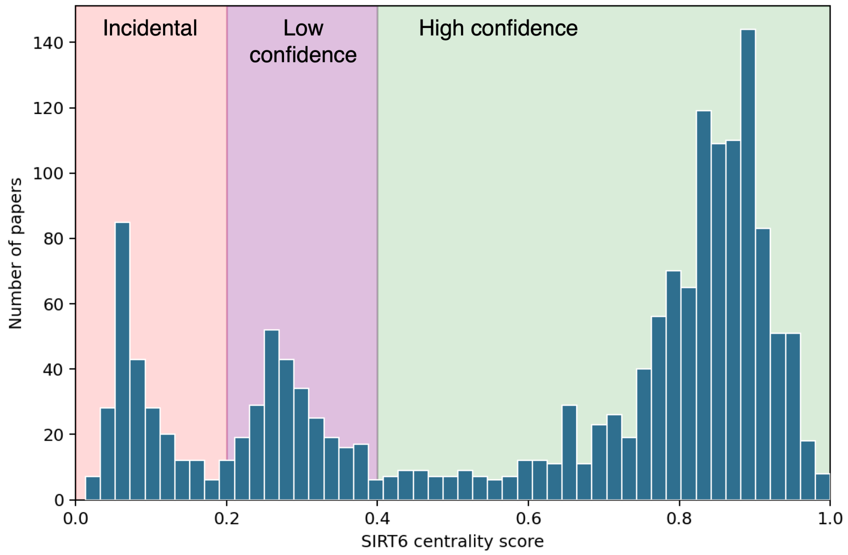


**Fig. 1** Topic modeling analysis of SIRT6-related papers. Histogram shows distribution of SIRT6 centrality scores, calculated for selected papers abstracts relevant for SIRT6 biology.

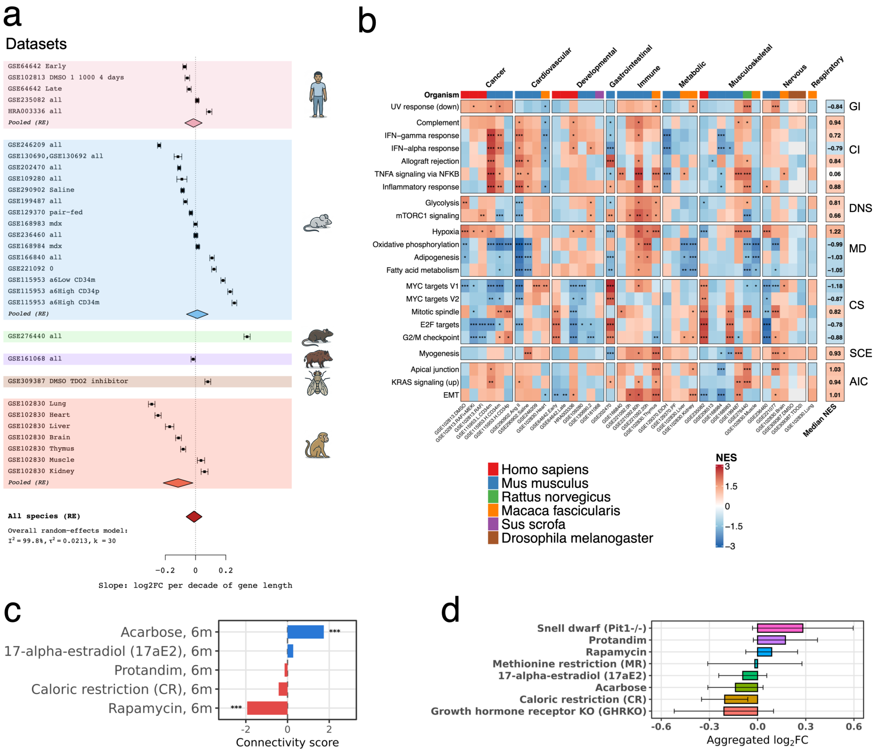


**Fig. 2** Functional changes in SIRT6-targeted models across species. (a) Forest plot showing the association between SIRT6 perturbation-induced expression changes and gene length, quantified as the slope of a weighted linear regression of $\log_2(\mathrm{FC})$ on $\log_{10}$ gene length. A negative slope indicates preferential downregulation of long genes, whereas a positive slope indicates preferential upregulation. (b) Heatmap illustrating the enrichment analysis of significant genes in SIRT6-perturbation datasets using MSigDB hallmark gene signatures. Datasets are stratified by a corresponding biological system and MSigDB hallmark terms classified by the relevance to aging hallmarks. "GI" denotes genomic instability, "CI" represents chronic inflammation, "DNS"- deregulated nutrient sensing, "MD"- mitochondrial dysfunction, "CS" - cellular senescence, "SCE" - stem cell exhaustion, and "AIC" represents altered intercellular communication hallmark. (c) Associations between transcriptomic signatures of longevity interventions and those observed in hepatic SIRT6-KO models. (d) Aggregated $\log_2(\mathrm{FC})$ values for SIRT6 expression in mouse liver following age-related interventions.

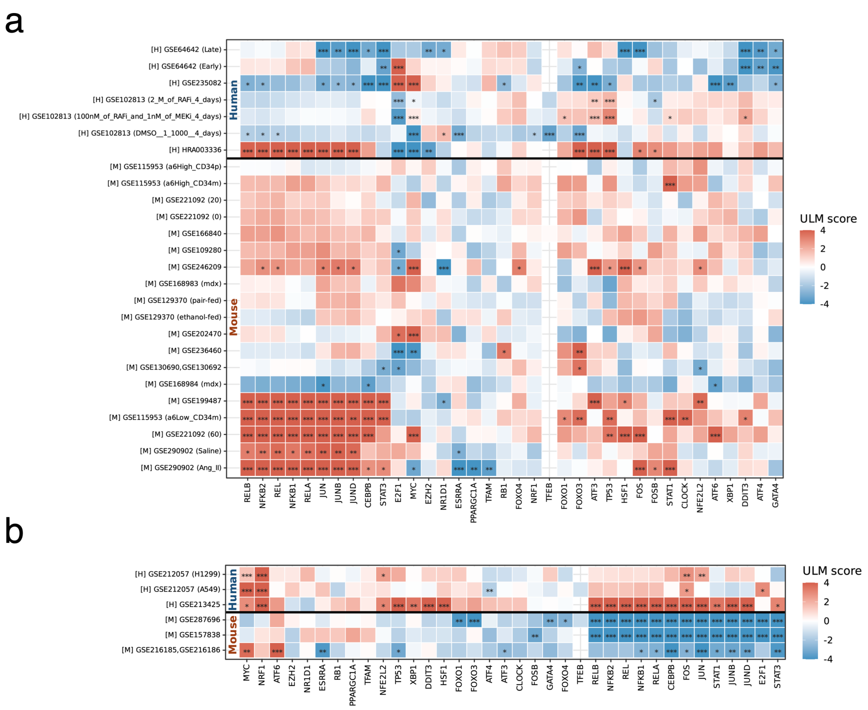


**Fig. 3** TF responses in mouse and human SIRT6 perturbation models. TF activity is shown for (a) SIRT6-knockout (KO) datasets and (b) SIRT6-overexpression datasets. In both panels, colors represent the $\Delta$ULM score. Asterisks indicate statistical significance: $***p < 0.001$, $**p < 0.01$, and $*p < 0.05$.

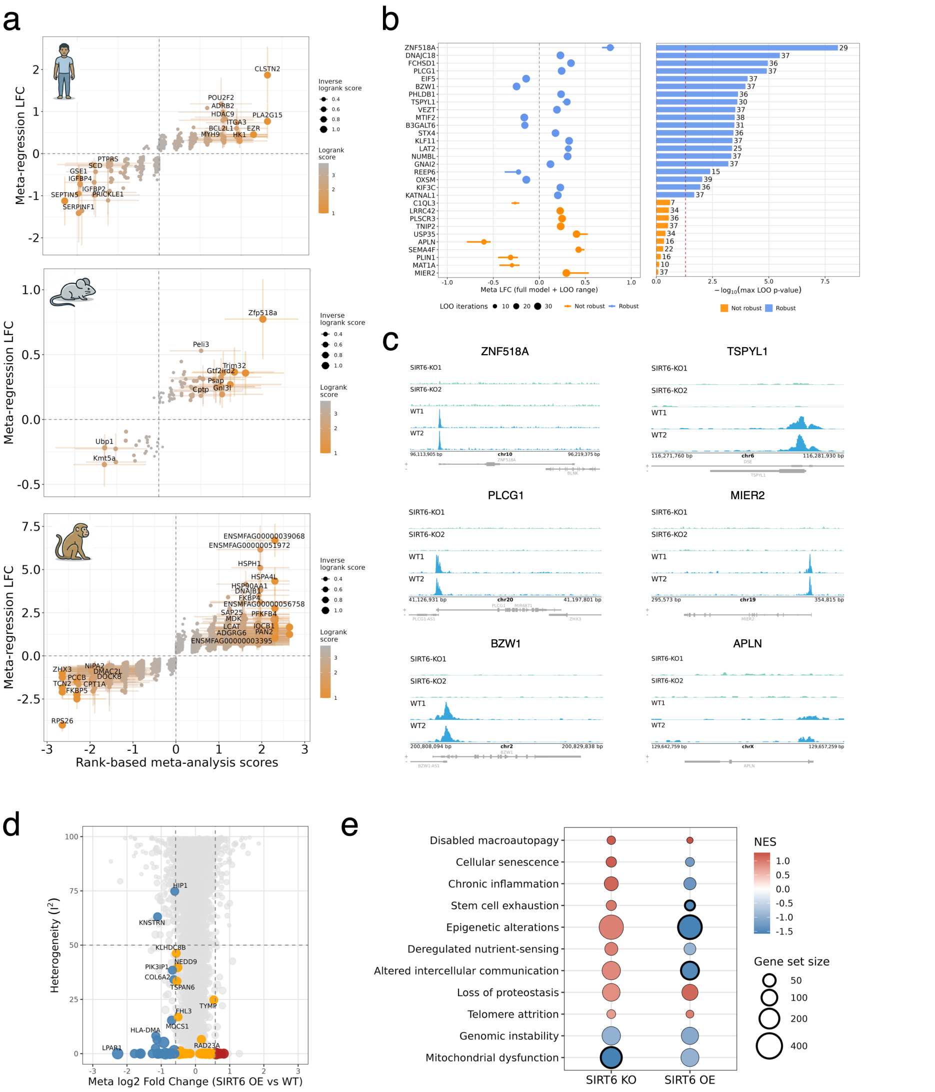


**Fig. 4** Meta-analysis of SIRT6-targeted experiments. (a) Scatter plot illustrating the most affected targets identified by two meta-analysis approaches in human (upper panel), mouse (middle panel) and macacue (lower panel) only. (b) Meta $\log_2$FC estimations after the leave-one-out procedure (left panel) and corresponding leave-one-out p-values (right panel), calcualted for significant genes. (c) CUT&RUN profiles of SIRT6 binding to the promoters of genes identified as signatures of SIRT6 deficiency. (d) Volcano plot showing the regression-based meta-analysis results for SIRT6-OE datasets across species. Highlighted circles shows genes, statistically significant in a meta-analysis (e) Dot plots representing enrichment values obtained from GSEA on aging gene signatures from Open Genes database for SIRT6 deficiency and overexpression experiments. Circled dots indicate FDR p-values $< 0.05$.

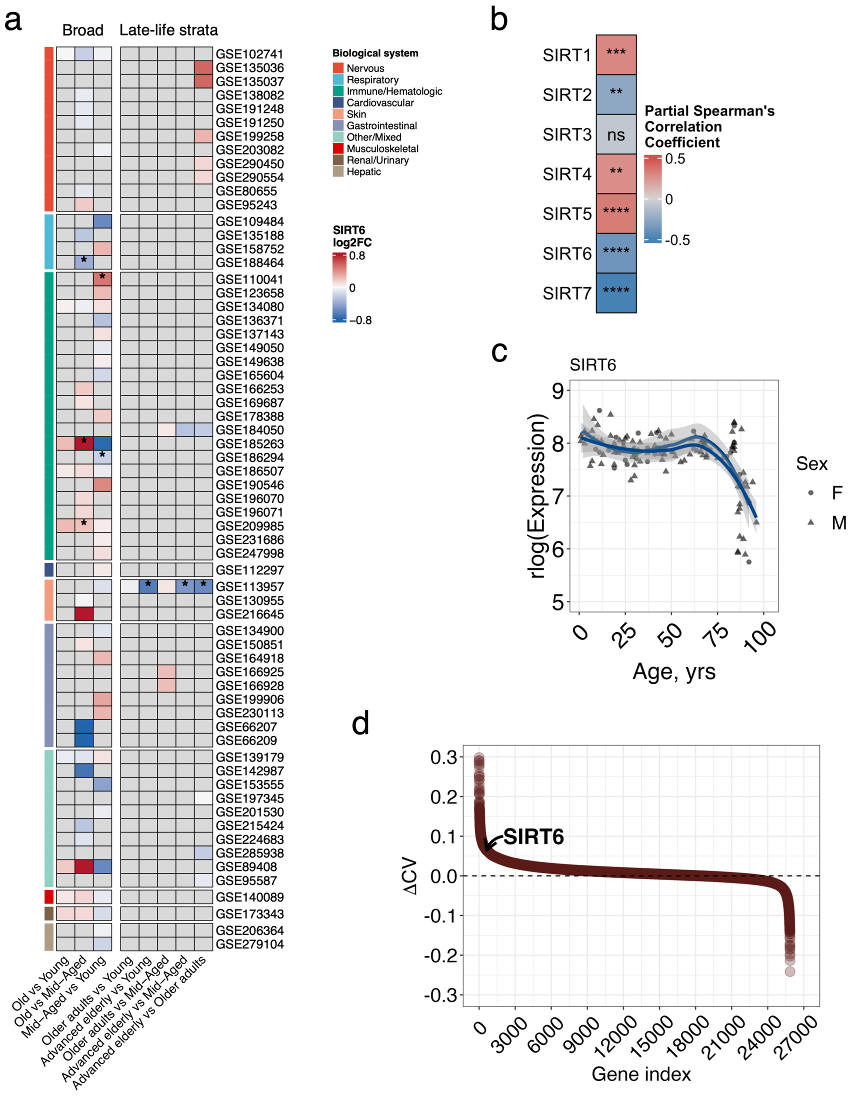


**Fig. 5** Transcriptional changes in SIRT6 during normal and pathological aging. (a) Heatmap of SIRT6 expression changes in the selected ARCHS4 datasets. Asterisks indicate FDR-adjusted $p < 0.05$. (b) Heatmap illustrating partial Spearman correlation of sirtuin expression with chronological age. (c) Age-related trajectory of SIRT6 expression in skin fibroblasts dataset (GSE113957) (d) Gene ranking by $\Delta$CV representing the expression variability in the oldest age range (from 72-84 to 84-96 years).
